# Density modification of cryo-EM maps

**DOI:** 10.1101/2020.05.13.091884

**Authors:** Thomas C. Terwilliger, Oleg V. Sobolev, Pavel V. Afonine, Paul D. Adams, Randy J. Read

**Affiliations:** New Mexico Consortium, Los Alamos NM 87544, USA; Bioscience Division, Los Alamos National Laboratory, Mail Stop M888, Los Alamos, NM, 87545, USA; Molecular Biophysics and Integrated Bioimaging Division, Lawrence Berkeley National Laboratory, Berkeley, CA, 94720, USA; Department of Bioengineering, University of California Berkeley, Berkeley, CA, USA; Cambridge Institute for Medical Research, Cambridge, CB2 0XY, UK

## Abstract

Density modification uses expectations about features of a map such as a flat solvent and expected distributions of density in the region of the macromolecule to improve individual Fourier terms representing the map. This process transfers information from one part of a map to another and can improve the accuracy of a map. Here the assumptions behind density modification for maps from electron cryomicroscopy are examined and a procedure is presented that allows incorporation of model-based information. Density modification works best in cases where unfiltered, unmasked maps with clear boundaries between macromolecule and solvent are visible and where there is substantial noise in the map, both in the region of the macromolecule and the solvent. It also is most effective if the characteristics of the map are relatively constant within regions of the macromolecule and the solvent. Model-based information can be used to improve density modification, but model bias can in principle occur. Here model bias is reduced by using ensemble models that allow estimation of model uncertainty. A test of model bias is presented suggesting that even if the expected density in a region of a map is specified incorrectly by using an incorrect model, the incorrect expectations do not strongly affect the final map.

**Synopsis:** The prerequisites for density modification of maps from electron cryomicroscopy are examined and a procedure for incorporating model-based information is presented.

## 1. Introduction

### 1.1. Construction and resolution-dependent accuracies of cryo-EM maps

Single-particle electron cryomicroscopy has recently become a powerful tool for high-resolution visualization of macromolecules. Nearly 400 three-dimensional electric potential maps of macromolecules (also referred to as density maps or just maps) have already been determined at resolutions of 3 Å or better and interpreted in terms of atomic models and deposited in the EM Data Bank and the Protein Data Bank (Berman *et al.*, 2000, Lawson *et al.*, 2011).

Three-dimensional cryo-EM maps are obtained by combining information from many two-dimensional images (Penczek, 2010a, Scheres, 2012, Nogales, 2016, Marques *et al.*, 2019). Each image is a projection of the macromolecule along the direction of the incident electron beam. The two-dimensional Fourier transform of an image containing one macromolecule corresponds to a central slice through the three-dimensional Fourier transform of the molecule. Using a set of such projections with sufficient coverage, the three-dimensional Fourier transform of the molecule can be sampled on a set of grid points with indices *hkl* that are very similar to crystallographic structure factors. This yields Fourier terms that can in turn be used to calculate a cryo-EM map representing the three-dimensional electric potential map for that macromolecule.

The accuracies of these Fourier terms can be estimated from a comparison of nominally independent determinations, each using half of the data. This comparison is known as the half-dataset Fourier Shell Correlation or FSC, (Rosenthal & Henderson, 2003). The FSC is typically very high (close to 1) at low resolution where the Fourier terms represent large-scale features and then decreases at higher resolution where the terms represent fine detail. The resolution where the FSC falls to 0.143 (and the estimated shell correlation between the map and true map falls to ½) is sometimes referred to as the “resolution” of the map (Rosenthal & Henderson, 2003). As cryo-EM maps are typically quite accurate at low resolution, the utility of a map is usually limited by the accuracies of the high-resolution Fourier terms and by appropriate weighting of low-resolution and high-resolution terms.

### 1.2. Improvements in obtaining accurate high-resolution cryo-EM maps

Many improvements in the construction and optimization of cryo-EM maps have been made in the past several years. Advances in the collection and analysis of images and in algorithms for using them to construct the three-dimensional Fourier transform of the macromolecule have led to dramatic improvements in the accuracies of high-resolution Fourier features in cryo-EM maps and to corresponding enhancements in the interpretability of the maps (Nogales, 2016). Additional advances have been made in optimizing the resolution-dependent weighting of Fourier terms to produce maps with improved clarity by map sharpening or blurring (Jakobi *et al.*, 2017, Terwilliger *et al.*, 2018, Ramlaul *et al.*, 2019, Ramírez-Aportela *et al.*, 2019).

### 1.3. Density modification

Recently efforts have been made to further improve the accuracy of cryo-EM maps by incorporation of prior knowledge about expected characteristics of these maps (Scheres, 2012, Kimanius *et al.*, 2020, Terwilliger, Ludtke, *et al.*, 2020), an approach sometimes known as “density modification”. This density modification approach is closely related to the procedure used in macromolecular crystallography with the same name (Wang, 1985, Podjarny *et al.*, 1996, Terwilliger, 2001a, Cowtan, 2010). The basic idea of density modification is that information about expected values of density in one part of a map can be used to partially correct errors in Fourier terms representing that map, and the corrected Fourier terms lead to an improved map everywhere (Terwilliger, 2001a). In this work, we will examine the assumptions about errors that allow density modification to work and show how model-based information can be incorporated into density modification of cryo-EM maps. We will focus on the form of density modification known as maximum-likelihood or statistical density modification (Terwilliger, 2001a), but other forms can have a similar interpretation.

### 1.4. Prerequisites for density modification to work

There are two key prerequisites for density modification to work. The first is that there are features or characteristics of the map that are known in advance. For example, it might be expected that a map of a macromolecule can be separated into two regions, one containing the macromolecule and having a distribution of density values similar to those found for other macromolecules, and the other containing the solvent and having nearly constant density. Alternatively, it might be known that some symmetry is present in the structure that was not used in the reconstruction process. In a case with unused symmetry, a target map could be constructed in which the target density in one part of the map is the average observed density in symmetry-related locations (Cowtan, 2000, Terwilliger, 2002, Cowtan, 2010). Each point in such a target map could have an associated uncertainty that describes the accuracy of the target map. This target map and its uncertainty could then be used to calculate a likelihood function that describes the plausibility of any map under consideration (Terwilliger, 2002).

The second prerequisite for density modification to work is that errors are local in Fourier space and global in real space. Specifying that errors are local in Fourier space means that the errors in any pair of Fourier terms with differing indices (*hkl*) are uncorrelated. If errors are local in Fourier space then they will be global in real space, and *vice versa* (Cowtan, 2000, Kammler, 2007). This second prerequisite is therefore equivalent to assuming that an error in one Fourier term (in Fourier space) will lead to correlated errors everywhere in the map (in real-space) and that the errors in different Fourier terms are independent. The effects of masking and of placing a small object in a large box on correlations of neighboring terms will be discussed below.

### 1.5. How density modification works and why these prerequisites are important

Density modification (either for cryo-EM or crystallography) can be thought of as a process with two basic steps. In the first step, one term (*hkl*) is removed from the Fourier transform of the map and all other terms are fixed. Continuing the first step, the value of the removed Fourier term that maximizes the plausibility (likelihood) of the map in the context of the other fixed terms is found. The resulting Fourier terms, called “map-phasing estimates (Terwilliger, 2001b), collectively represent a new estimate of the Fourier transform of the map.

In the second step, an improved estimate of each Fourier term is obtained from a weighted average or recombination of the original and map-phasing estimates, and these weighted average Fourier terms are used to calculate a final density modified map. The details of this second step vary considerably and involve half-datasets in cryo-EM but not in crystallography, but it always involves recombination of original and map-phasing estimates of Fourier terms.

The first step in this process requires some way of calculating the likelihood or plausibility of a map. This leads to the first prerequisite of having some features of the map that are known in advance. Note that expected features of the map can be specified in a probabilistic fashion and different probability distributions can be used in different places in a map (Terwilliger, 2001a).

The second (recombination) step will work best if the map-phasing estimates of the Fourier terms have errors that are uncorrelated with the errors in the original Fourier terms. This is because averaging works best if errors are not correlated. As described below, this leads to the second prerequisite, that errors must be local in Fourier space.

### 1.6. Why errors must be local in Fourier space to obtain independent map-phasing estimates of Fourier terms

To see why having errors that are local in Fourier space will lead to map-phasing estimates of Fourier terms that have errors uncorrelated with errors in the original Fourier terms, consider first a special case. Suppose we have a situation where all Fourier terms except one are known perfectly, where the one Fourier term that has some error is the one we are interested in, and where the value of the density is known to be zero in some “solvent” region of the map.

The error in the one Fourier term causes errors throughout the map, including the solvent region. The map-phasing value of the one Fourier term of interest will simply be the value that yields zero in the solvent region of the map. Since the rest of the Fourier terms are perfect and our knowledge of the value in the solvent region is exact, this will yield the exact value of the one Fourier term we were unsure about. As a result of this operation, the density in the entire map will become perfect. Note that in this process, information has been transferred from one part of the map (knowledge of the value of density in a small region) to another part of the map (the rest of the map becomes perfect).

The reason this transfer of information is possible is that there is redundant information present. The complete set of Fourier terms is by itself sufficient to define the perfect map. Taking out one Fourier term removes information, but knowledge of the flat solvent region is more than sufficient to define the value of that Fourier term. Another way to look at this is to note that the one unknown Fourier term in this example is correlated with (and in this case, defined by) all other Fourier terms through an interference or G-function (Rossmann & Blow, 1962, Abrahams & Leslie, 1996, Kleywegt & Read, 1997) that comes from the flat solvent region.

In the idealized situation above, there were no errors in the fixed Fourier terms or in the expectations about the map. In a realistic case, there will be errors involved, and whether these errors are correlated will influence the utility of density modification. The error in a map-phasing estimate of a particular Fourier term will come from errors in the other (fixed) terms and from errors in specification of expectations about the map. The key concept is that if errors in the Fourier space representation of the map are local, then the errors in the fixed terms will be independent of the error in the Fourier term that was removed, and therefore the error in the map-phasing estimate of this Fourier term will be independent of the error in its original value. This means that if errors are local in Fourier space, information from the original map can be combined in a simple way with information from map-phasing to yield an improved density-modified map.

There is an important opposite case to note as well. Suppose that the starting map in the example above was systematically incorrect, for example from instrumental errors leading to large noise peaks or by inclusion of an incorrect model at some stage. All the Fourier terms would have errors that reflect this systematic error in the map and the errors in the Fourier terms for this systematically incorrect map would not be local. If one Fourier term were removed and its value chosen based on a flat solvent, an incorrect value would be obtained.

In addition to the requirement of local errors in Fourier space, for map phasing to work optimally, errors in specification of the expectations about the map must be independent of the errors in the map itself. An extreme case where this would not be true, and density modification would not work at all, would be one in which prior knowledge about a map is simply a target map with values equal to the original map. The opposite and highly useful case would be one in which prior knowledge about a map is a target map with errors that are completely independent of the errors in the starting map. For example, a reconstruction might be carried out in which some local molecular symmetry is not exact and is therefore not included in the averaging process. In such a case, a target map for one copy of this molecule could be generated from another (independent) copy of this molecule.

### 1.7. The effect of masking (or finite support) on correlations in Fourier space

If any point in a map can in principle have any value, then no correlations are expected between nearby terms in Fourier space. However, if there are volumes in the map that have a constant value (which can trivially be set to zero just by changing the F_000_ term in Fourier space), the map can be said to have finite support as there is only a limited volume that has non-zero values. In that case, there will be correlations between nearby Fourier terms.

If a map is masked (set to zero or some other fixed value outside of the masked region of the map), then neighbouring Fourier terms will necessarily be correlated because the map will now have finite support, even if it did not before masking. The fall-off of correlations with separation in Fourier space depends on how much of the map is masked (Kammler, 2007, Penczek, 2010b), because the correlations are determined by the Fourier transform of the mask, termed the G-function in crystallography (Rossmann & Blow, 1962, Abrahams & Leslie, 1996). At one extreme, if no mask is applied to a map lacking finite support then the values of all Fourier terms are uncorrelated. At the other extreme, if the region inside the mask is infinitely small and at the origin of the cell or grid, then all Fourier terms are identical.

### 1.8. Correlations in Fourier terms for the true map

The true (but unknown) map is expected to possess non-random features (e.g., flat regions, bounded values or non-random distributions of values, symmetry), so the true Fourier terms that represent this map will be correlated. These correlations among the true Fourier terms make phase improvement possible in macromolecular crystallography (Rossmann & Blow, 1962, Bricogne, 1974, Cowtan, 2000). Also, as described in section 1.6, these correlations among the true Fourier terms are the basis for obtaining new estimates of a Fourier term by map phasing.

### 1.9. Potential for correlations of errors in Fourier terms due to masking

The observed map can be thought of as a sum of a weighted true map and a noise map. If the values in the noise map are all drawn from the same distribution and are independent of position, then the error terms they contribute to the observed Fourier terms will have no local correlations.

It is common in the construction of cryo-EM maps to include procedures that either explicitly or implicitly lead to masking of the final map (Tang *et al.*, 2007). This masking leads to correlations in the errors of neighbouring Fourier terms and reduces the effectiveness of density modification. Note that the masking typically improves the starting map so that the optimal procedure is not necessarily obvious (Terwilliger, Ludtke, *et al.*, 2020).

### 1.10. Rationale for assumption of errors that are local in Fourier space

A justification of the assumption of local errors in Fourier space comes from the way that three-dimensional Fourier terms are estimated from the two-dimensional Fourier transforms of individual images. The value of the Fourier transform of the molecule at to a particular index (*hkl*) is estimated from the subset of central sections (Fourier transforms of individual images) that overlap with that location (*hkl*) in the Fourier transform (Penczek, 2010a). These subsets of central sections will generally be different for each location (*hkl*) in the transform. As each central section comes from a separate image, with different errors in the original image and in its orientation, the errors in different central sections and therefore errors at different locations in Fourier space (*hkl*) are likely to be largely independent. The extent to which this assumption holds is examined in section 3.1.

### 1.11. Probability distribution for errors in Fourier terms

We expect that the assumption of independent errors will apply to Fourier terms with the same indices *hkl* coming from different images. Since the Fourier terms in the final reconstruction result from averaging all of these observations, the central limit theorem leads us to expect that the true and observed Fourier terms will be related by the complex normal distribution (Scheres, 2012, Terwilliger, Ludtke, *et al.*, 2020). The consequences of this are discussed in section 2.2 below.

## 2. Methods

### 2.1. Apoferritin maps and model

We used high-resolution (1.8 Å, EMD-20026) and low-resolution (3.1 Å, EMD-20028) maps of human apoferritin available from the EMDB (Lawson *et al.*, 2011, Pintilie *et al.*, 2020). The crystal structure of human apoferritin (Masuda *et al.*, 2010) was docked into and refined against the 1.8 Å cryo-EM map with the *Phenix* tools *dock_in_map* (Liebschner *et al.*, 2019) and *real_space_refine* (Afonine, Poon, et al., 2018).

### 2.2. Estimation of complex errors in Fourier terms for human apoferritin

The (complex) errors in Fourier terms for human apoferritin were estimated using a comparison of Fourier terms representing high-resolution (1.8 Å) and low-resolution (3.1 Å) maps of human apoferritin. For the low-resolution map (EMD-20028) we used just one half-map in this analysis in order to have a dataset with large errors. Based on the half-dataset FSC for the high-resolution map, we previously concluded that the Fourier terms representing this map are very accurate up to a resolution of at least 3 Å, with estimated correlation to Fourier terms representing the true map of about 0.98 up to this resolution (Terwilliger, Ludtke, *et al.*, 2020). We therefore used the Fourier terms for the high-resolution map as an estimate of those representing the true map.

As discussed above, we assume that the errors in the reconstructions are drawn from a complex normal distribution (Terwilliger, Ludtke, *et al.*, 2020). As a consequence, the differences between the high- and low-resolution reconstructions also can be expected to be drawn from a complex normal distribution. As the high-resolution reconstruction is by far the more accurate of the two, the errors are expected to be dominated by the errors in the low-resolution reconstruction.

A complex normal distribution relating two complex numbers (such as the Fourier terms from cryo-EM reconstructions) can be described by the complex correlation between them. For a resolution shell, the complex correlation *CC* is equivalent to the FSC value for that shell, as shown in (1).

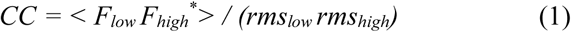

Where *F*_*low*_ is a Fourier term for the low-resolution map, *F_high_*^*^ is the complex conjugate of the corresponding term for the high-resolution map, and *rms_low_* and *rms_high_* are the rms values of terms for the low- and high-resolution maps, respectively. In crystallography, the parameter *σ*_A_ is analogous to the FSC, as it is also a complex correlation coefficient (Srinivasan & Chandrasekaran, 1966) describing the relationship between the true structure factor and the model structure factor. The expected value of one Fourier term when the other is known is given in (2).

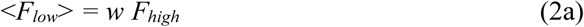

where

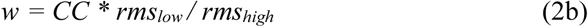

The error (*E*) in the low-resolution estimate (*F*_*low*_) of a Fourier term can be estimated from the difference between low- and high-resolution estimates of that term,

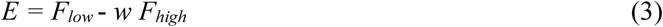

Note that the error (*E*) is assumed here to be independent of *F*_*high*_. The weight *w* is analogous to the Luzzati *D-*factor (Luzzati, 1952) in the crystallographic difference map coefficient, *mF_O_ – DF_C_* (Read, 1986): *D* bears the same relationship to *σ*_A_ as *w* does to *CC*.

To estimate the error in each low-resolution Fourier term of apoferritin, we calculated Fourier terms representing the high- and low-resolution apoferritin maps to a resolution of 3 Å. Then in shells of resolution, the rms value of terms for each map (*rms_high_, rms_low_*) and the Fourier shell correlation (*CC*) between Fourier terms for each map was obtained. The estimated complex error (*E*) in a particular Fourier term was then calculated by weighted subtraction of the high-resolution Fourier term (*F*_*high*_) from the low-resolution Fourier term (*F*_*low*_), as given in (3) above.

### 2.3. Estimation of local variation in a map

Estimates of the local standard deviation of values in a map were obtained in shells surrounding the center of the map. The value reported for a shell from radius *r_1_* to radius *r_2_* is the standard deviation of the values at all grid points with distance from the origin in the range of *r_1_* to *r_2_*.

### 2.4. Generation of model-based density and its local uncertainty using ensemble models

Ensemble models representing a macromolecule were created with two different procedures. In each procedure, the maps used to create a model were density-modified maps calculated without model information. Also in each case, two groups of models were created, one refined using one of the density-modified half-maps for that macromolecule and one refined against the other. By default, 8 models were created in each group.

The first procedure for generation of one model was to start from a supplied model, apply a shaking procedure to the model, and carry out real-space refinement of the partially-randomized model using one density-modified half map as a target. This procedure is carried out using the *Phenix* (Liebschner *et al.*, 2019) tool *mia*. An additional 7 models were then created using the same procedure and the same half-map but different random seeds, and an additional 8 using the other half map.

The second procedure for generation of a single model was to use the *Phenix* tool *map_to_model* to automatically build a model based on a full density-modified map, then to refine that model based on one density-modified half map as a target. Then additional models were generated by running *map_to_model* over again with a different seed used in all steps that involve randomization and varying the procedure used in density modification and model-building. The variation in density modification consisted of all four possible combinations turning on and off the density modification parameters *blur_by_resolution* and *spectral_scaling* (Terwilliger, Ludtke, *et al.*, 2020). These are two parameters that a user might normally vary if density modification does not work well. The *blur_by_resolution* parameter applies a blurring B-value to the final map where the B-value is given by 10 times the resolution (Terwilliger, Ludtke, *et al.*, 2020). The *spectral_scaling* parameter rescales the final map to have a resolution dependence similar to that of a model protein in a box of solvent (Terwilliger, Ludtke, *et al.*, 2020). The variation in model-building procedure by default simply consists of turning the parameter *high_density_from_model* on and off when running the *Phenix* tool *map_to_model*. This parameter is normally varied in this way in a default run of *map_to_model* as well (Terwilliger, Adams, *et al.*, 2020).

Once an ensemble of models was created based on one density-modified half-map, this ensemble was used along with the corresponding half-map to create an ensemble-based density target and associated uncertainty. First, all the grid points in the map that were within at least (by default) 3 Å of an atom in at least 3 of the members of the ensemble were identified. These represent the grid points where model-based information can be obtained along with some estimate of its uncertainty. Then at all such grid points, the mean and standard deviation of density were calculated. The local uncertainty in density values was taken to be the local standard deviation of density, after smoothing with a radius by default equal to 2/3 the resolution.

This ensemble-based density for one half map was then directly combined with the corresponding density-modified half map. The purpose of this step is to generate as accurate a target map as possible. We note that including the density-modified map directly in this procedure can result in some correlation of errors in the target map with errors in the starting map. The combination of model and density-modified maps was carried out with weights based on the uncertainties in each map. The local uncertainty in the density-modified half map was estimated by calculating the smoothed squared difference between the two density-modified half maps, using a smoothing radius (by default) equal to the resolution of the original map. The value of the target map at each point was then simply the weighted average of the ensemble model-based map and the density-modified half map. Finally, the target map was masked to include only those grid points where at least (by default) three models had contributed to the estimate of ensemble-based density.

## 2.5. Density modification including information represented as a target map with local uncertainties

We used *RESOLVE* maximum-likelihood density modification (Terwilliger, 2000, 2001b) to carry out density modification including model-based information represented as a target map with local uncertainties. The procedure is very similar to that previously used for density modification of a single cryo-EM half map (Terwilliger, Ludtke, *et al.*, 2020). The grid representing the map was divided into the region where model-based information was present (inside the masked portion of the target map described in the previous section), and where it was not (outside the mask). In the region where model-based information was present, the target map was used along with the corresponding uncertainty map as the basis for creating a log-likelihood function for each grid point. Elsewhere, the log-likelihood function was the same one that would be used in density modification without a model (Terwilliger, 2000, 2001b). The log-likelihood of any set of map values was calculated by summing the log-likelihood values at all grid points.

### 2.6. Software availability

The density modification tool *phenix.resolve_cryo_em* is available as part of the *Phenix* software suite (Liebschner *et al.*, 2019).

## 3. Results and Discussion

### 3.1. Evaluating the assumption that cryo-EM errors are local in Fourier space

A central assumption in density modification is that errors are local in Fourier space and global in real space. This assumption is important because (cf. section 1.6) it is the independence of errors in Fourier space that allows useful recombination of information based on expectations about a map (map phasing) with the original information present in a map. We examine this assumption here using a comparison of Fourier terms representing a high-resolution map (1.8 Å) and a lower-resolution map (3.1 Å) of human apoferritin.

Our strategy was to calculate the (complex) error in each Fourier term representing the low-resolution map by subtraction of weighted terms from the high-resolution map (assumed to be very accurate in this resolution range; see Methods). The Fourier transform of these error terms then shows where the errors lie in real space. If errors are local in Fourier space then the errors in real space are expected to be uniformly distributed throughout the map. If instead errors are local in real space, then the errors in real space might be confined to some part of the map. In particular, they might be confined to the region where the protein is located or at least might be much larger in that region.

Fig. 1 shows slices through a model of human apoferritin and through cryo-EM maps of apoferritin. Fig 1A shows a slice through the model and illustrates that apoferritin is essentially a spherical shell of protein. Fig. 1B shows the same slice through the center of the high-resolution (1.8 Å) map, calculated after truncation of Fourier terms at a resolution of 3 Å and weighting Fourier terms using Eq. (2b). As expected, this slice through the map shows a clear ring containing protein-like density. Fig. 1C shows the low-resolution (3.1 Å) half-map of apoferritin. This map is similar to the high-resolution map but has considerable density outside the protein region. Note that the density in this slice of the map does not extend quite all the way to the edge of the map, indicating that the map has been masked outside of this region. Finally, Fig. 1D shows the estimated error in the low-resolution map, given using Eq. (3) as the difference between the maps in Fig. 1C and Fig. 1B. The high density in this estimated error map is distributed across all parts of the map, with somewhat more high density near the center of the map and less density far from the center. In particular, note that the level of density in the estimated error map in the region where the protein is located is not substantially different than nearby regions outside the protein.

**Figure 1.**
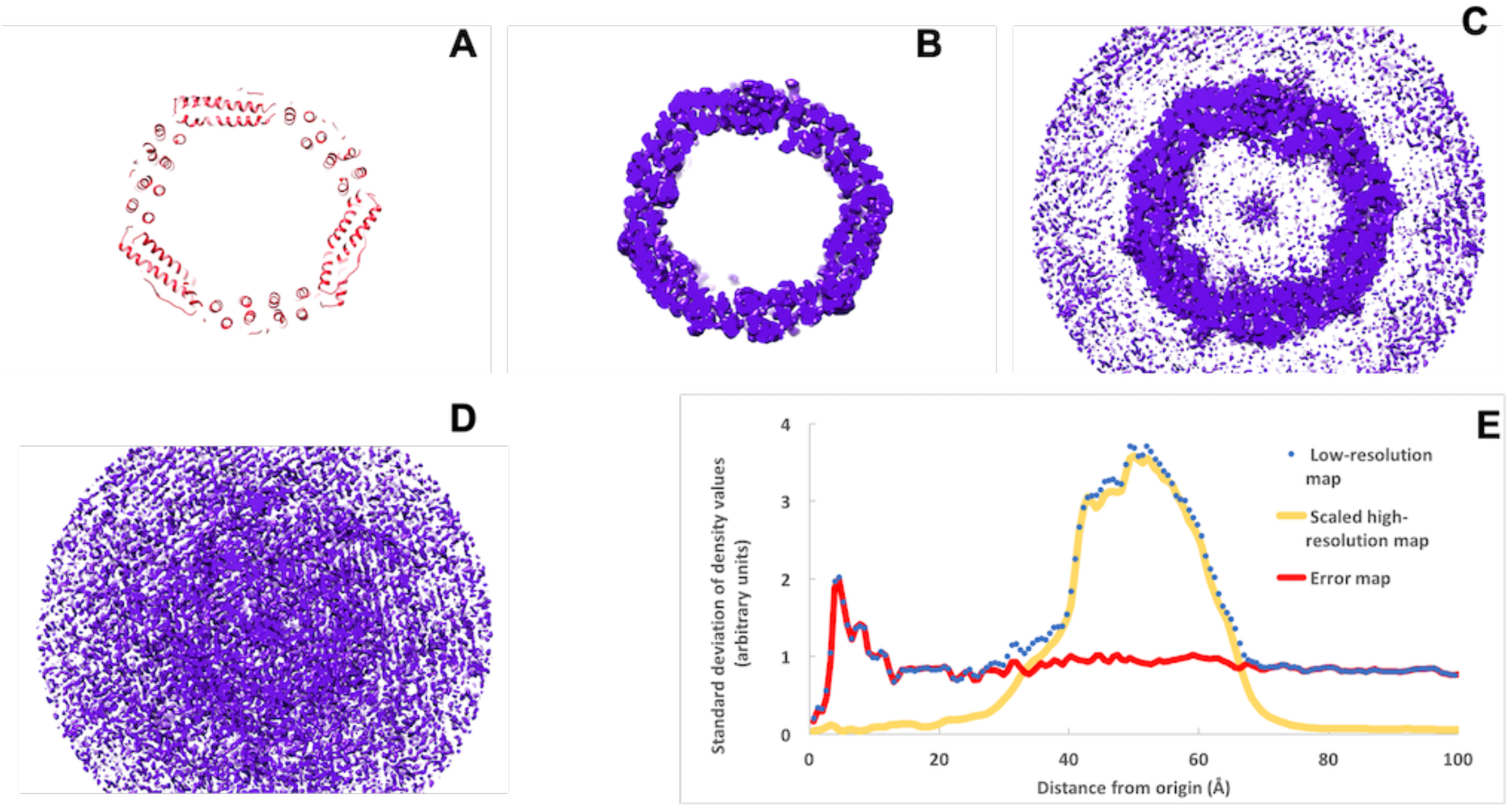
Sections through a model and cryo-EM maps of human apoferritin. A. Section through model of human apoferritin. B. Section as in A, through cryo-EM map (EMD 20026) at resolution of 1.8 Å, scaled as described in the text. C. Section as in B through half-map (EMD 20028) at resolution of 3.1 Å. D. Difference between the maps in C and B. E. Local standard deviation of the maps in B (yellow, labeled scaled high-resolution map), C (blue dots, labeled low-resolution map), and D (red, labelled error map) in radial shells around the centers of the maps. The contour levels in B and C are at 0.65 times the rms of the maps and in D is at 1.4 times the rms of the map. The scale in E is arbitrary but the same for all three maps. All maps drawn with Chimera (Pettersen *et al.*, 2004).

Fig. 1E shows the local standard deviations (local variation) of the three maps shown in Figs. 1B, 1C and 1D, plotted as a function of distance from the center of the map. The local standard deviation of density values for the scaled high-resolution map (from Fig 1B) is very small (about 0.1 on an arbitrary scale) except in a range of radii from about 40 Å to 60 Å, where the values range from 2 to 4. The local standard deviations of density values for the low-resolution map (from Fig. 1C) are very similar to those in the high-resolution map in the range of radii from about 40 Å to 60 Å. Outside of that range, the standard deviations of density values of the low-resolution map are reduced (values of about 1), but still much higher than those of the high-resolution map (values of about 0.1). Finally, the local standard deviation of values in the estimated error map (from Fig. 1D) is relatively constant over all radii from zero to 100 Å, with values of about 1. Taking Fig. 1D and Fig. 1E together, this indicates that at least in this case, errors are not localized in the region where the protein is located. This result is consistent with the hypothesis that errors are global in real space and localized in Fourier space.

### 3.2. Including additional sources of information in density modification of cryo-EM maps

A central element in density modification is the use of prior knowledge about likely features or characteristics of the map. In our previous work on density modification of cryo-EM maps, this prior knowledge included expectations of a flat solvent region and expectations that distributions of density values in the protein region would be similar to those seen previously or obtained after averaging half-maps (Terwilliger, Ludtke, *et al.*, 2020). In crystallographic density modification, several other important sources of prior knowledge have been used. Here we will focus on using density based on an ensemble-model representation of a structure as a source of prior information about a map (Perrakis *et al.*, 1999, Terwilliger *et al.*, 2007, Herzik *et al.*, 2019). For comparison and as a simpler example, we will start by considering the expectation that density in the true map may have symmetry that is not imposed (Bricogne, 1974, Wang, 1985, Terwilliger, 2002, Cowtan, 2010).

For clarity in this discussion about additional sources of information about a map, we will frame prior knowledge as a target map with uncertainties that may vary from one part of the map to another (Terwilliger, 2001a), though it could take other forms including any kind of map likelihood function (Terwilliger, 2000). In this simple framework, non-crystallographic symmetry can be represented as values in the target map in one region that are derived from a different but symmetry-related region in the map (Terwilliger, 2002). Uncertainties in symmetry can be derived from variances among symmetry-related regions. Similarly, model-based density derived from a single model could be included in a target map with an overall uncertainty. Alternatively, averaged densities from multiple models could be included with uncertainties derived from the variation among the models. As in our previous work, information about expected distributions of densities in the region of the macromolecule and solvent can also be included (Terwilliger, Ludtke, *et al.*, 2020).

### 3.3. Considerations in specification of prior knowledge about a map

There are three aspects of the specification of prior knowledge that can affect the utility of density modification. One is whether the errors in the target map are correlated with the errors in the original map. As discussed above (cf. section 1.6), the target map errors must be independent of the errors in the original map to be most useful. A second is whether the errors in the target map are systematically biased in a way that could lead to misinterpretation of the final map, an effect known as model bias (Ramachandran & Srinivasan, 1961). This could in principle occur, for example, if the target map is derived from a single model. Incorrectly-placed atoms in that model could lead to density in the target map that show those atoms, and if that information were propagated through the density modification procedure the final map might show density for those atoms even though they are not present. A third is whether the uncertainties in the target map are accurately specified. The uncertainties need to be reasonably accurate to optimally weight information from the target map and original map in the recombination step of density modification.

In the case of using symmetry present in a map that was not imposed during reconstruction, these three aspects of the specification of prior knowledge are relatively simple. The errors in the target map are essentially independent of those in the original map because they come from a different part of the map. The errors in the target map are generally not biased in a way that would cause misinterpretation, and the uncertainties in specification can be estimated relatively accurately by comparison of all the symmetry-related copies. The principal limitation of this approach is that the symmetry-related copies may not in fact be identical. This limitation can be addressed at least in part by allowing local variability by estimating the uncertainty in the target map locally (Abrahams & Leslie, 1996, Terwilliger, 2002).

In contrast, when model-derived information is used, estimation of errors is challenging and model bias is an important consideration. Errors in the target map could be correlated with those in the original map because the model or models are generally derived from that original map. Note that this correlation of errors might be very high if every detail of the original density is reflected in the model, or quite small if there are many constraints used in creating the model so that the details of the model are not coming from the original density. If part of the model-derived target map is incorrect, the density in the target map might still look like a macromolecule, leading to the possibility of misinterpretation (Ramachandran & Srinivasan, 1961). Further, if a single model is used to generate a target map, estimation of the uncertainty in that target map may be difficult.

### 3.4. Density modification with information based on ensemble models

To address two of the challenges in using model-based information for density modification, we use target density derived from an ensemble-model representation of a structure. The basic idea is to generate a group of models that are as diverse as possible while maintaining compatibility of each with experimental data and prior structural knowledge. Then the variation among these models represents a lower limit on the actual uncertainty in any one model (Terwilliger *et al.*, 2007, Herzik *et al.*, 2019). In the context of creating a target map for density modification, the target density at each point in the map is the average density calculated from members of the ensemble, with uncertainty estimated from its standard deviation (see Methods).

This straightforward estimation of uncertainties in the model-based target map is an important reason for using ensemble-model representation. Additionally, this approach indirectly reduces the chance of incorrect model-based density looking like an actual feature of a macromolecule. The ensemble-model approach uses density averaged over a group of plausible models. If the map is unclear in a certain region and constraints on the models do not restrict their conformations in that region, the ensemble of models generated for that region will normally have a wide variety of configurations, leading to high estimated uncertainty and lack of clear features in the model-based map. Consequently, incorrectly-modeled regions will typically have few features using this approach, while a single-model representation would look just like a macromolecule, but would be incorrect.

The ensemble-model representation does not in itself address the issue of correlation of errors in the model-based map with errors in the original map. As noted above, however, this correlation can be reduced to low values by inclusion of restraints and constraints in the generation of each model in the ensemble (i.e., by not overfitting the models).

### 3.5. Application of density modification with ensemble models to apoferritin maps

We tested the utility and limitations in model-based density modification of cryo-EM maps by applying it to maps of human apoferritin at resolutions of 1.8 Å and 3.1 Å. In each case two half-maps were used as a starting point. These maps were first density-modified without model-based information. Then an ensemble-model representation of each corresponding half-map was created by independently building 8 models using the *Phenix* tool *map_to_model* (Terwilliger, Adams, *et al.*, 2020) and used in density modification.

Fig. 2 shows regions of these low-resolution (3.1 Å) apoferritin maps along with a reference model refined using a density-modified version of the high-resolution map. Fig. 2A shows the original 3.1 Å apoferritin map. This map has been sharpened using half-map sharpening in which the resolution-dependent scale on the map is adjusted to yield RMS values of Fourier terms that are proportional to values of the Fourier shell correlation in each shell of resolution (Terwilliger *et al.*, 2018). Fig. 2B shows the map obtained using density modification but no model-based information. Fig. 2C shows the map obtained using an ensemble model for apoferritin based on models generated automatically with the *Phenix* tool *map_to_model*. Comparing the density at locations indicated by arrows in Figs. 3A, 3B, and 3C, it can be seen that the density-modified map shows an increase in detail over the original map, and the map density-modified with an ensemble model shows considerable detail not present in either of the others.

**Figure 2.**
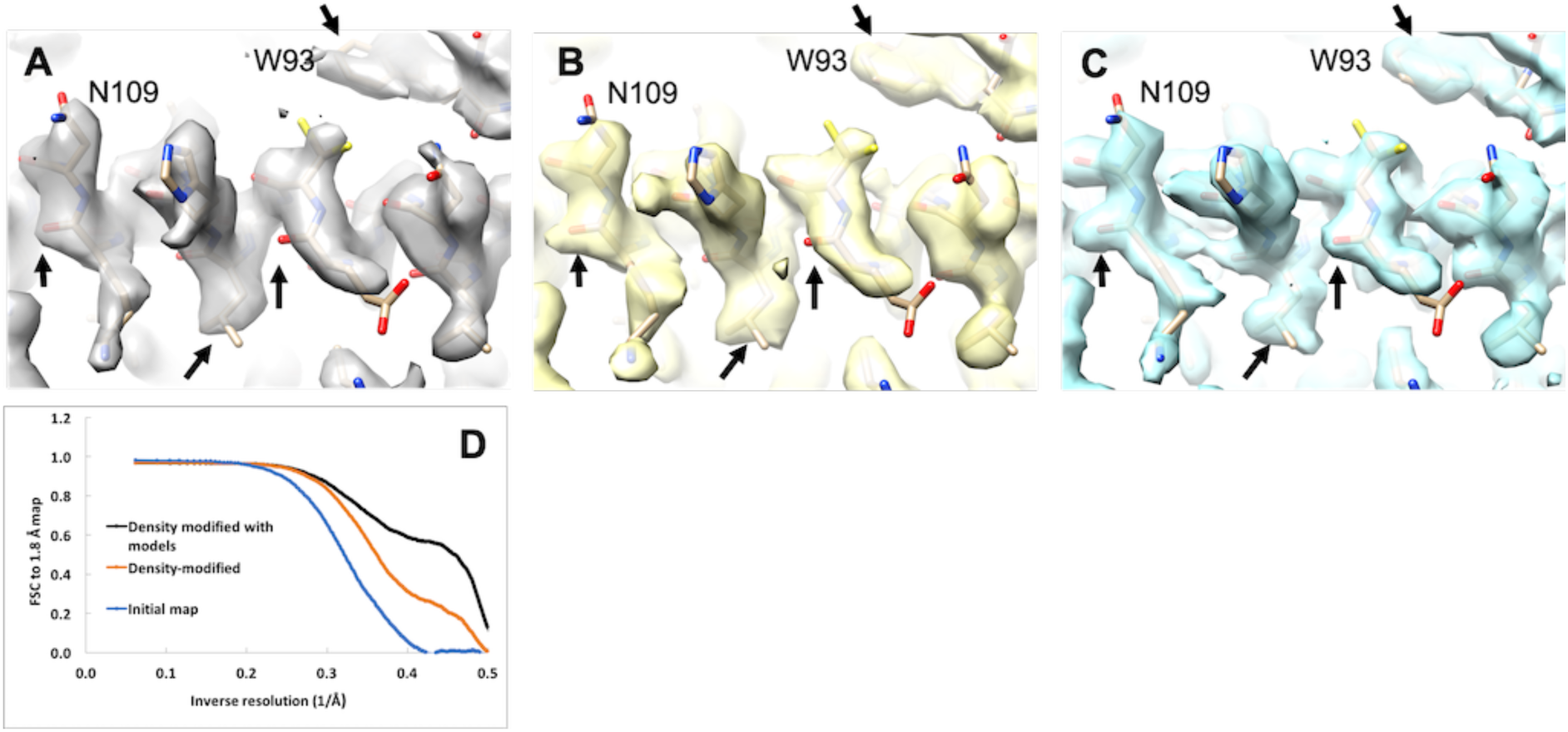
Low-resolution (3.1 Å) apoferritin maps. A. Original 3.1 Å apoferritin map, half-map sharpened (Terwilliger *et al.*, 2018). B. Map obtained using density modification but no model-based information. C. map obtained using an automatically-generated ensemble model. The model shown is from the crystallographic structure of human apoferritin (Masuda *et al.*, 2010), re-refined against the 1.8 Å apoferritin map (EMD-20026). Arrows indicate locations in the map that where the clarity is improved by density modification. D. Fourier shell correlation of the maps shown in A (blue, labelled Initial map), B (orange, labelled Density-modified) and C (black, labelled Density modified with models) with the high-resolution (1.8 Å) apoferritin map (EMD 20026). The density modification analysis was carried out to a resolution of 2 Å so all Fourier terms from the density modified map are zero beyond that resolution and therefore all FSC values are zero as well.

**Figure 3.**
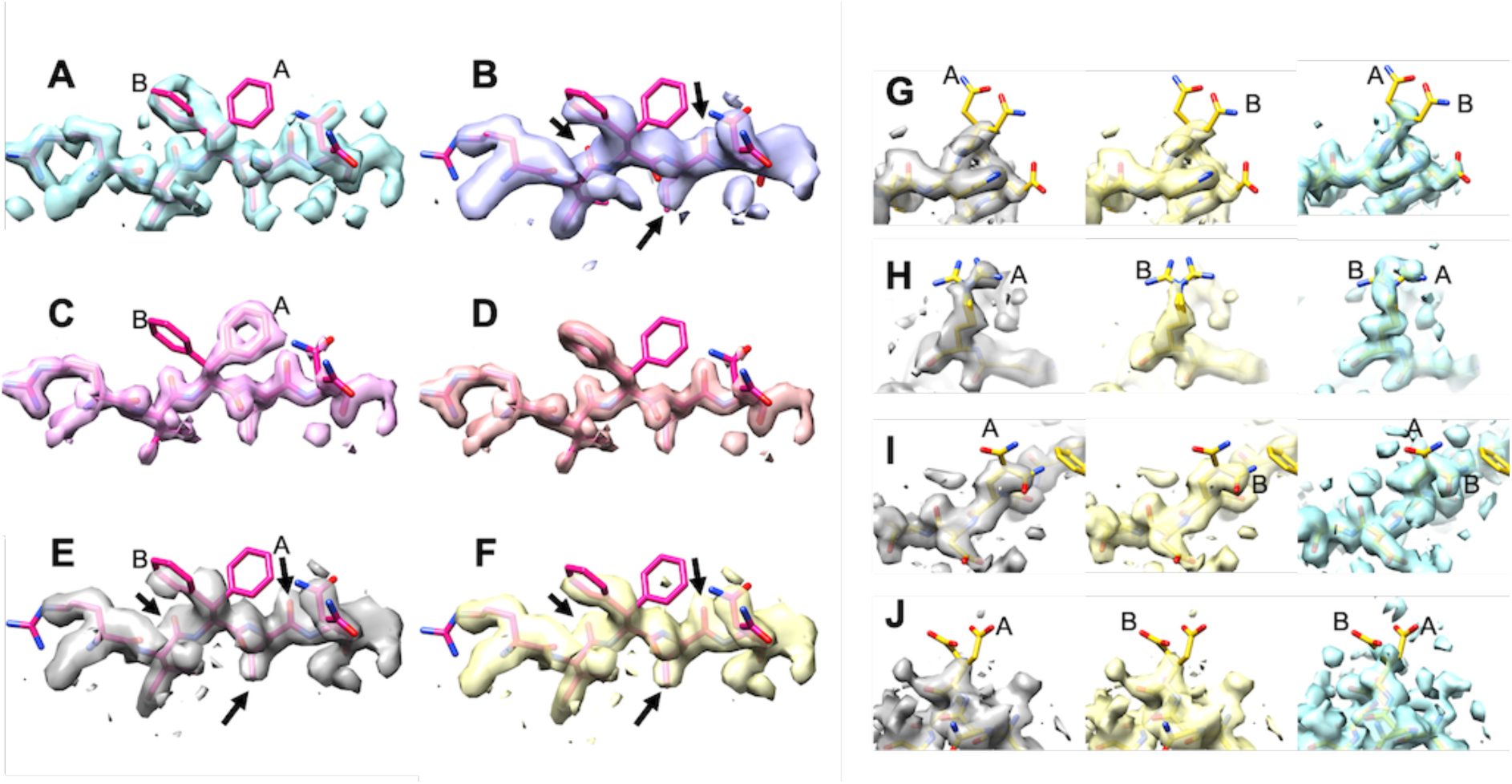
Control analysis using model density based on incorrect side-chain conformations. Panels A-F are analyses varying target density for F81 and G-J are varying target density for residues Q14, R22, Q83 and E140. A. High-resolution (1.8 Å) apoferritin map (EMD 20026), density modified without model information. A crystal structure of human apoferritin, docked into and partially refined against the EMD 20026 map, but still containing multiple conformations of several side chains is shown. B. Low-resolution (3.1 Å) apoferritin map (EMD 20028), density modified without model information. C. Target density using conformation “A” of F81. D. Target density using conformation “B” of F81. E. Density-modified 3.1 Å apoferritin map including target density from conformation “A” shown in panel C. F. Density-modified 3.1 Å apoferritin map including target density from conformation “B” shown in panel D. Panels G-J each show three maps focused on a selected residue (Q14 in G, R22 in H, Q83 in I and E140 in J). The maps on the left and middle in each case are density-modified 3.1 Å apoferritin maps including target density from conformation “A” or “B” of the selected residue, respectively. The map on the right is the corresponding high-resolution (1.8 Å) density-modified apoferritin map. “A” and “B” side chain conformations are indicated. Contours for members of each pair of density-modified maps are set at identical levels.

Fig. 2D illustrates the improvement in the density modified maps estimated by Fourier shell correlation of each map with the high-resolution (1.8 Å) map of apoferritin. It can be seen that at a resolution of 2.5 Å (inverse resolution of 0.4 Å^−1^), the original low-resolution map has a very low correlation (0.05) with the high-resolution map, while the density-modified map has a higher correlation of about 0.3 and the map density-modified with an ensemble model has a correlation of about 0.6.

As test of model bias, single-model representations of apoferritin were created in which one side chain was deliberately placed in a correct or incorrect conformation. Then the resulting model was used in density modification without further adjustment. Fig. 3 (panels A-F) show the results of this control analysis in which a model with a partially correct or incorrect side chain at residue F81 is used in density modification. Fig. 3A shows the high-resolution apoferritin map (1.8 Å, EMD 20026), density modified without model information, and showing a docked and partially refined crystal structure of human apoferritin (Masuda *et al.*, 2010) as a reference. The high-resolution density map shows a conformation of F81 similar to that of the “B” conformer in this model. Fig. 3B shows the map using lower-resolution (3.1 Å) information (EMD 20028), also density modified without model information, and also indicating that the “B” conformer is correct. figs. 3C and 3D show the target density generated from the models with conformations “A” or “B” of F81, respectively. As expected, each of these shows clear density for the corresponding conformation of F81. Finally, figs. 3E and 3F show the density modified maps using target density from Fig. 3C or 3D. Fig. 3E shows that despite target density identifying conformation “A” of F81, the density-modified map shows density for conformation “B”. Fig. 3F, as expected, also shows density for conformation “B”. Note that in the map density-modified using model information in Fig. 3E, details indicated with black arrows are clearer than in the map density modified without model information shown in Fig. 3B. This suggests that even though some aspects of the model information used in Fig. 3E are incorrect (i.e., the side chain of F81 is in the wrong conformation), other aspects of the model information can still improve corresponding parts of the map.

It is possible that the lack of model bias shown in Fig. 3A-F is due to the very clear density for the side chain of Y81 in the low-resolution map density-modified without a model. Panels G-J repeat this control analysis at four additional residues with varying clarity of side chain density. In each panel, the left and middle images show maps density-modified using model-based information with either the “A” or “B” conformer of a selected side chain. The right image in each panel shows the corresponding region of the high-resolution (1.8 Å, EMD 20026) density-modified map as a reference.

Panel G shows this analysis for residue Q14. In this example neither the high-resolution map nor the model-based density-modified maps show significant density for either conformation of the side chain, despite each using target density for one conformation in the process.

Panel H shows the analysis for residue R22. The side-chain density in the high-resolution map on the right does not show clear density for a single conformation, but it appears somewhat more consistent with the “A” conformation shown of R22 than the “B” conformation. The map density-modified using the “A” conformation target density shows some density for the “A” conformation, while the one using the “B” conformation does not have significant side-chain density. As the correct conformations are not known in this case, interpretation of this result is not straightforward. The result appears similar to that found for Y81 above, however, in which correct information coming from a model may improve corresponding parts of a map while incorrect information has a relatively small effect.

Panel I illustrates an analysis for residue Q83. In this case the high-resolution map shows the “B” conformation of Q83 is present. Density modification using either conformation “A” or “B” as a target yields a map showing conformation “B”. This result is similar to the one for residue Y81 in panels A-F.

Finally, panel J (E140) shows a case similar to the case for Q14 in panel G in which neither conformation is clear in the high-resolution map on the right. Also similar to the case for Q14, neither model-based density modified map shows substantial side chain density for either conformation of the side chain.

In this control experiment, we created artificial target density that in some cases was completely incorrect and in other cases was partially or largely correct. In no cases did the density-modified maps show any obvious sign of the incorrect conformations. At the same time, the density-modified maps are generally improved in locations away from incorrectly-modeled side chains.

There are two reasons why this is possible. One reason is that the information coming from the original map (naturally) shows the side chains in their correct conformations (see the map density-modified without model information in panel B). This means that the incorrect model information would have to have a strong effect on the final density in order to show the incorrect conformation.

The second and perhaps more important reason this result is possible is that the map-phasing step in density modification is not simply adjusting Fourier terms to yield a map matching target density. This would indeed show the incorrect conformation present in the target density. Instead, each new Fourier term is one which, in the context of all the other (original) Fourier terms, yields a map that best matches overall target density. This distinction is crucial because all the other Fourier terms contain information about the true density in the map. In the context of the other Fourier terms, the value of one Fourier term that leads to a map showing the incorrect side chain density would (in general) lead to a map that disagrees with other expectations about the map. Consequently, if most of the information (e.g., target map, solvent and macromolecule density distributions) used to evaluate map plausibility is correct, even serious localized errors in expectations about the map may not lead to substantial errors in the density-modified map.

### 3.6. Application of density modification with ensemble models to maps at resolutions from 3.4 to 4.4 Å

We carried out density modification with automatic generation of ensemble models using half-maps from six cryo-EM maps with resolutions from 3.4 Å to 4.4 Å. Fig. 4 illustrates the maps obtained for two of these cases. In each case three maps are shown: the initial map, sharpened automatically using model-based sharpening with the *Phenix* tool *auto_sharpen* (Terwilliger *et al.*, 2018), the standard density-modified map, and the density modified map obtained using ensemble models. Figs. 4A, 4B, and 4C show these maps for the human polycystin-2/PKD2 TRP channel (EMDB 8200 and PDB entry 5k47), at a resolution of 4.2 Å (Grieben *et al.*, 2017). The density modified map in panel 4B shows additional detail not present in the original map in Fig. 4A (note the side chain for F605), and the map density modified using ensemble models shows further detail (note the side chain for M603). Figs. 4D, 4E and 4F show the same series of maps for the human CLC-1 chloride ion channel (EMDB 7544 and PDB entry 6coy) at a resolution of 3.4 Å (Park & MacKinnon, 2018). The map density modified using ensemble models again shows the greatest degree of clarity (note the density corresponding to hydroxyl oxygens and the side chain of L361 marked with arrows in the three maps).

**Figure 4.**
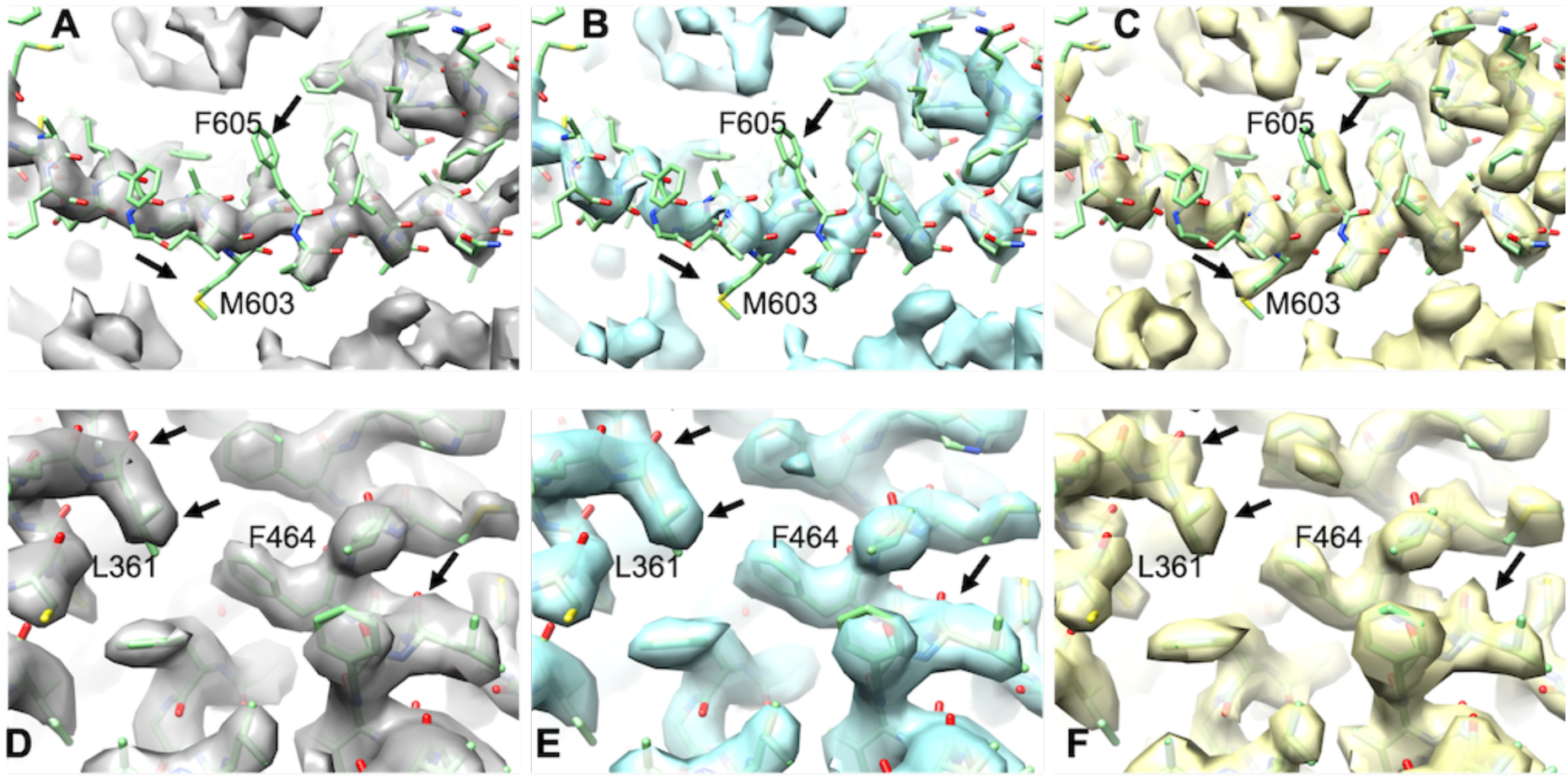
Comparison of original maps (A, D), density modified maps (B, E), and maps density modified using automatically-generated ensemble models (C, F). A-C. Maps for the human polycystin-2/PKD2 TRP channel at a resolution of 4.2 Å (EMDB 8200 and PDB entry 5k47). D-F. Maps for the human CLC-1 chloride ion channel at a resolution of 3.4 Å (EMDB 7544 and PDB entry 6coy). Differences between maps are noted with arrows and selected residues in the deposited models are labeled. Maps in each set are contoured to enclose equal volumes (Urzhumtsev *et al.*, 2014).

Fig. 5 illustrates the map improvement for each of the 6 maps we examined as measured by the correlation between the density each map and masked density calculated from the corresponding deposited model. These map correlations were calculated up to the resolution at which density modification was carried out in order to capture the range of resolution where improvement was expected. These resolutions were, on average, 0.8 Å finer than the nominal resolution of these maps. It can be seen from Fig. 5 that in each case the map correlation for the map that had been density-modified using information from ensemble models was higher than that for either the corresponding original or standard density-modified maps. The improvements are small (average improvement in map correlation of 0.02) but may be useful in map interpretation (cf. Fig. 4). Note that when map-model correlations were calculated only up to the nominal resolution of the maps, the model-based density modified maps were slightly poorer than the original maps (average difference in map correlation of −0.02), indicating that the procedure for recombination of original and map-phasing maps may not be fully optimal.

**Figure 5.**
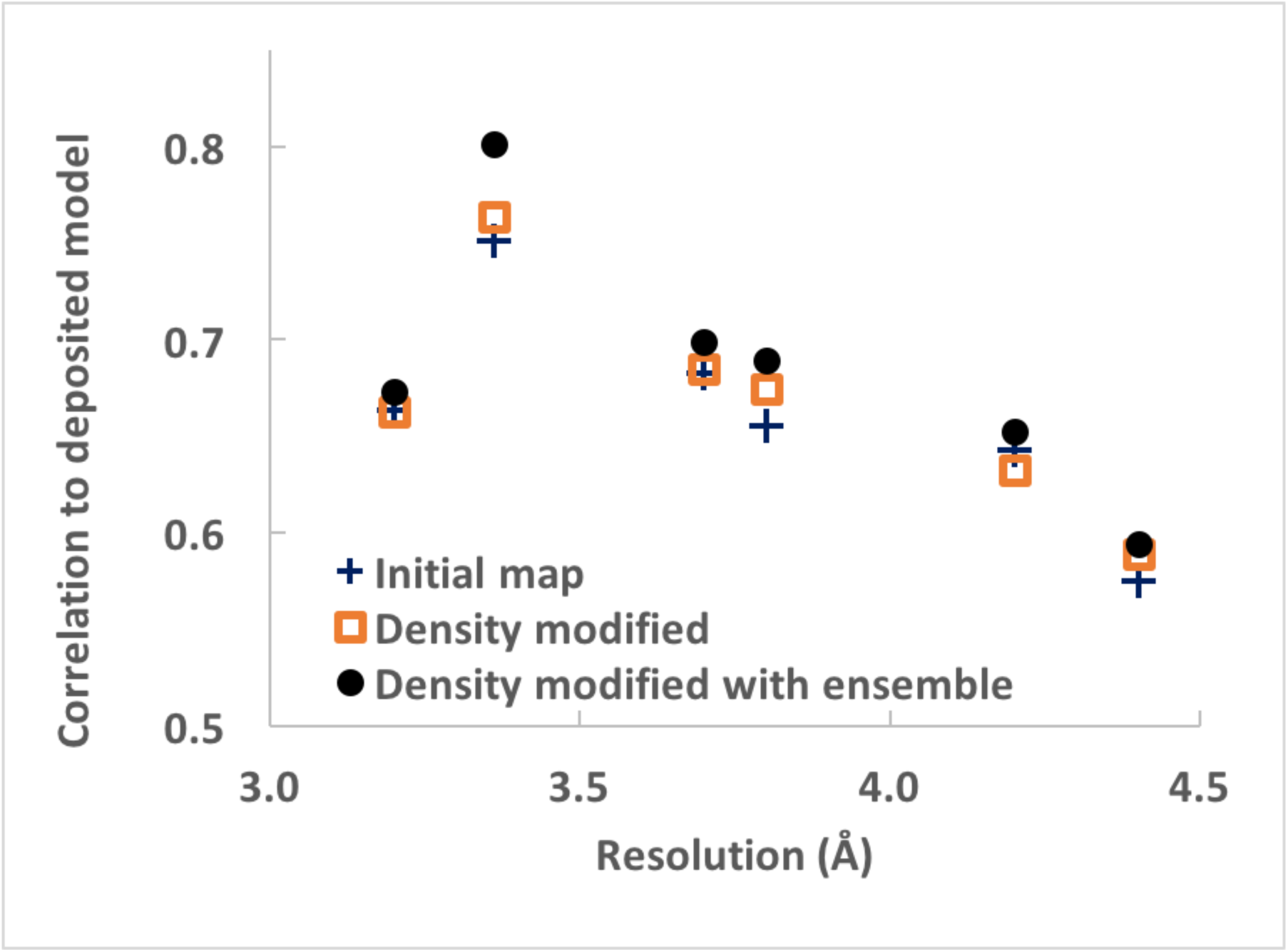
Map improvement using density modification and using density modification with ensemble models. Six cryo-EM maps were density modified as in Fig. 4 and the correlation between each map and model-based density was calculated as described in the text. The maps and models were EMDB 7573 and PDB entry 6crv, 3.2 Å (Kirchdoerfer *et al.*, 2018); EMDB 7544 and PDB entry 6coy, 3.4 Å (Park & MacKinnon, 2018); EMDB 3837 and PDB entry 5ood, 3.7 Å (Merino *et al.*, 2018); EMDB 3805 and PDB entry 5ogw, 3.8 Å (Pospich *et al.*, 2017); EMDB 8200 and PDB entry 5k47, 4.2 Å (Grieben *et al.*, 2017); EMDB 7582 and PDB entry 6cs2, 4.4 Å (Kirchdoerfer *et al.*, 2018). Map-model correlations for the initial maps are shown with a blue plus sign, those for the density-modified maps are shown with orange squares, and those for maps density modified with ensemble models are shown with black circles. Map-model correlations (*CC_mask_*) were obtained out with the *Phenix* tool *map_model_cc* (Afonine, Klaholz, *et al.*, 2018, Liebschner *et al.*, 2019). Correlations were obtained using deposited models after setting the atomic displacement parameters (B-values) of all the atoms in the models to zero, a procedure shown previously to yield a useful metric of map quality (Terwilliger *et al.*, 2018). This procedure also has the advantage here of removing bias in the model favoring the deposited map. The resolution used for map-model correlations are the resolutions used in density modification.

### 3.7. Limitations

Most of the limitations of density modification of cryo-EM maps are related to the prerequisites described in the introduction. The first prerequisite was that there is substantial prior knowledge about characteristics of the map available. Limitations that relate to this prerequisite include:

(1) Density modification will not work well if the solvent and protein regions are specified incorrectly. This could happen if the map is very noisy and the solvent boundary is not placed correctly. It could also occur if the expected volume occupied by the macromolecule is not appropriate, as might happen for example if the macromolecule includes substantial surrounding lipid.

(2) It could also work poorly if the uncertainties about the expected values of the map are systematically over- or under-estimated.

(3) If constant values for density in the solvent region are assumed, density modification could fail if there is substantial variation in this region. This might occur if lipid is present and is included in the solvent region. It can also occur if the solvent region includes the entire map and the density variation falls off far away from the macromolecule.

(4) Procedures such as ours involve cutting out the region containing the macromolecule and some solvent around it, and using this boxed region as is, not placing it in a larger box. This reduces the variation in expected density values in the solvent region. As a side effect, however, this boxing cuts out some of the information about errors in Fourier terms that was reflected in the solvent region that was removed. Additionally, as the boxing is normally done automatically, it can potentially cut out part of the macromolecule, leading to a loss of information. Boxing procedures can also lead substantial truncation errors manifested as high noise levels at the edges of the box.

(5) If the characteristics of the macromolecule vary substantially from one location to another and this is not included in the expectations about the map, density modification could work sub-optimally. This can often occur as cryo-EM maps frequently have regions with differing effective resolutions. Focused reconstructions in which one region of the map is much higher resolution than other parts could work poorly for this reason as well.

(6) If the two-half maps used in the analysis share systematic errors, then the error analysis used to recombine density-modified and original maps will not be accurate and density modification may not work optimally.

The second prerequisite for density modification is that errors are local in Fourier space and therefore global in real space. This prerequisite can fail to be satisfied in several ways:

(1) If the maps are substantially masked or a small part of each maps is cut out and placed into a large box, density modification could work poorly because substantial correlations are introduced into Fourier terms by boxing and masking. Further, in this case the information about what is wrong with the Fourier terms has been cut out (for example the noise in the solvent region has information about the errors in Fourier terms). Note that this case is different than the boxing done in the present procedure where the box includes the macromolecule and surrounding solvent and the box of density is not placed in a larger empty box.

(2) If the maps have had local modifications such as local resolution filtering, then correlations of errors will have been introduced in these maps and density modification may work less well.

(3) If multiple cycles of density modification are carried out and Fourier terms are adjusted on the basis of other already-adjusted terms, correlations of errors can be introduced between Fourier terms.

(4) If fixed model density is used as a target with strong weighting and multiple cycles of density are carried out then the map-phasing map could end up looking like the model density. This effect can also be thought of as introducing correlated errors in the Fourier terms that specifically affect the density in the region of the macromolecule.

There are additional limitations of the method as well:

(1) Our implementation assumes that the two half-maps used in density modification have similar distributions of errors. If this is not the case then the weighting of half-maps could be sub-optimal.

(2) Another implementation-specific limitation is that it is assumed that errors are generally proportionally greater at high resolution than at low resolution.

(3) A general limitation is that maps can only be improved in the resolution ranges where the starting maps are poor. As most maps have relatively accurate low-resolution terms, this means that density modification may improve only a narrow high-resolution range of Fourier terms.

(4) In principle, inclusion of model-based information for one part of a map could improve the density everywhere, just as information about a flat solvent can improve density in the region of the macromolecule. In practice, we have not observed this. Rather, the density in the region where the model is specified appears to improve in locations where the model is correct and not to change compared to standard density modification in locations where the model is incorrect. Presumably the reason that information from models does not improve the map as much as knowledge about the solvent region is that the model-based information is not nearly as accurate as information about the flat solvent.

### 3.8. Conclusions

Density modification can be a useful procedure for improving the accuracy of cryo-EM maps. It is carried out starting with two half-maps and produces a density-modified final map. Density modification requires expectations about the density in the true map and requires that errors in the starting map are local in Fourier space. It works best in cases where unfiltered, unmasked maps with clear boundaries between macromolecule and solvent are visible and where there is substantial noise in the map, both in the region of the macromolecule and the solvent. In our present implementation, it also is most effective if the characteristics of the map are relatively constant within regions of the macromolecule and the solvent.

Model information can be included in density modification. Models can be supplied from other sources or can be automatically generated, and they can considerably improve density modification if the models are accurate. Model bias can in principle occur, but tests suggest that even if the expected density in a region of a map is specified incorrectly by using an incorrect model, the incorrect expectations do not strongly affect the final map. In our implementation, model bias is also reduced by construction of ensemble models that allow estimation of uncertainties.

## Acknowledgements

The authors appreciate support received from the US National Institutes of Health (grant P01GM063210 to P.D.A., R.J.R. and T.C.T.). This work was partially supported by the US Department of Energy under contract DE-AC02-05CH11231. R.J.R. is supported by a Principal Research Fellowship from the Wellcome Trust (grant 209407/Z/17/Z).

## References

Abrahams, J. P. & Leslie, A. G. (1996). Acta Crystallogr D Biol Crystallogr 52, 30–42.

Afonine, P. V., Klaholz, B. P., Moriarty, N. W., Poon, B. K., Sobolev, O. V., Terwilliger, T. C., Adams, P. D. & Urzhumtsev, A. (2018). Acta Crystallogr D Struct Biol 74, 814–840.

Afonine, P. V., Poon, B. K., Read, R. J., Sobolev, O. V., Terwilliger, T. C., Urzhumtsev, A. & Adams, P. D. (2018). Acta Crystallographica Section D 74, 531–544.

Berman, H. M., Westbrook, J., Feng, Z., Gilliland, G., Bhat, T. N., Weissig, H., Shindyalov, I. N. & Bourne, P. E. (2000). Nucleic Acids Res 28, 235–242.

Bricogne, G. (1974). Acta Crystallographica Section A 30, 395–405.

Cowtan, K. (2000). Acta Crystallographica Section D 56, 1612–1621.

Cowtan, K. (2010). Acta Crystallographica Section D 66, 470–478.

Grieben, M., Pike, A. C. W., Shintre, C. A., Venturi, E., El-Ajouz, S., Tessitore, A., Shrestha, L., Mukhopadhyay, S., Mahajan, P., Chalk, R., Burgess-Brown, N. A., Sitsapesan, R., Huiskonen, J. T. & Carpenter, E. P. (2017). Nat Struct Mol Biol 24, 114–122.

Herzik, M. A., Jr., Fraser, J. S. & Lander, G. C. (2019). Structure 27, 344–358.e343.

Jakobi, A. J., Wilmanns, M. & Sachse, C. (2017). Elife 6.

Kammler, D. W. (2007).

Kimanius, D., Zickert, G., Nakane, T., Adler, J., Lunz, S., Schönlieb, C.-B., Öktem, O. & Scheres, S. H. W. (2020). bioRxiv, 2020.2003.2025.007914.

Kirchdoerfer, R. N., Wang, N., Pallesen, J., Wrapp, D., Turner, H. L., Cottrell, C. A., Corbett, K. S., Graham, B. S., McLellan, J. S. & Ward, A. B. (2018). Sci Rep 8, 15701.

Kleywegt, G. J. & Read, R. J. (1997). Structure 5, 1557–1569.

Lawson, C. L., Baker, M. L., Best, C., Bi, C., Dougherty, M., Feng, P., van Ginkel, G., Devkota, B., Lagerstedt, I., Ludtke, S. J., Newman, R. H., Oldfield, T. J., Rees, I., Sahni, G., Sala, R., Velankar, S., Warren, J., Westbrook, J. D., Henrick, K., Kleywegt, G. J., Berman, H. M. & Chiu, W. (2011). Nucleic Acids Res 39, D456–D464.

Liebschner, D., Afonine, P. V., Baker, M. L., Bunkoczi, G., Chen, V. B., Croll, T. I., Hintze, B., Hung, L.-W., Jain, S., McCoy, A. J., Moriarty, N. W., Oeffner, R. D., Poon, B. K., Prisant, M. G., Read, R. J., Richardson, J. S., Richardson, D. C., Sammito, M. D., Sobolev, O. V., Stockwell, D. H., Terwilliger, T. C., Urzhumtsev, A. G., Videau, L. L., Williams, C. J. & Adams, P. D. (2019). Acta Crystallographica Section D 75, 861–877.

Luzzati, V. (1952). Acta Crystallographica 5, 802–810.

Marques, M. A., Purdy, M. D. & Yeager, M. (2019). Current Opinion in Structural Biology 58, 214–223.

Masuda, T., Goto, F., Yoshihara, T. & Mikami, B. (2010). Biochemical and Biophysical Research Communications 400, 94–99.

Merino, F., Pospich, S., Funk, J., Wagner, T., Küllmer, F., Arndt, H.-D., Bieling, P. & Raunser, S. (2018). Nat Struct Mol Biol 25, 528–537.

Nogales, E. (2016). Nat Methods 13, 24–27.

Park, E. & MacKinnon, R. (2018). eLife 7, e36629.

Penczek, P. A. (2010a). Methods in enzymology 482, 1–33.

Penczek, P. A. (2010b). Methods in Enzymology, edited by G. J. Jensen, pp. 73–100: Academic Press.

Perrakis, A., Morris, R. & Lamzin, V. S. (1999). Nat Struct Biol 6, 458–463.

Pettersen, E. F., Goddard, T. D., Huang, C. C., Couch, G. S., Greenblatt, D. M., Meng, E. C. & Ferrin, T. E. (2004). J Comput Chem 25, 1605–1612.

Pintilie, G., Zhang, K., Su, Z., Li, S., Schmid, M. F. & Chiu, W. (2020). Nature methods 17, 328–334.

Podjarny, A. D., Rees, B. & Urzhumtsev, A. G. (1996). Crystallographic Methods and Protocols, edited by C. Jones, B. Mulloy & M. R. Sanderson, pp. 205–226. Totowa, NJ: Humana Press.

Pospich, S., Kumpula, E.-P., von der Ecken, J., Vahokoski, J., Kursula, I. & Raunser, S. (2017). Proceedings of the National Academy of Sciences 114, 10636–10641.

Ramachandran, G. N. & Srinivasan, R. (1961). Nature 190, 159–161.

Ramírez-Aportela, E., Vilas, J. L., Glukhova, A., Melero, R., Conesa, P., Martínez, M., Maluenda, D., Mota, J., Jiménez, A., Vargas, J., Marabini, R., Sexton, P. M., Carazo, J. M. & Sorzano, C. O. S. (2019). Bioinformatics 36, 765–772.

Ramlaul, K., Palmer, C. M. & Aylett, C. H. S. (2019). Journal of Structural Biology 205, 30–40.

Read, R. (1986). Acta Crystallographica Section A 42, 140–149.

Rosenthal, P. B. & Henderson, R. (2003). J Mol Biol 333, 721–745.

Rossmann, M. G. & Blow, D. M. (1962). Acta Crystallographica 15, 24–31.

Scheres, S. H. (2012). J Mol Biol 415, 406–418.

Srinivasan, R. & Chandrasekaran, R. (1966). Indian J Pure Appl. Phys. 4, 178–186.

Tang, G., Peng, L., Baldwin, P. R., Mann, D. S., Jiang, W., Rees, I. & Ludtke, S. J. (2007). Journal of Structural Biology 157, 38–46.

Terwilliger, T. (2000). Acta Crystallographica Section D 56, 965–972.

Terwilliger, T. (2001a). Acta Crystallographica Section D 57, 1755–1762.

Terwilliger, T. (2001b). Acta Crystallographica Section D 57, 1763–1775.

Terwilliger, T. (2002). Acta Crystallographica Section D 58, 2082–2086.

Terwilliger, T. C., Adams, P. D., Afonine, P. V. & Sobolev, O. V. (2020). Protein Sci 29, 87–99.

Terwilliger, T. C., Grosse-Kunstleve, R. W., Afonine, P. V., Adams, P. D., Moriarty, N. W., Zwart, P., Read, R. J., Turk, D. & Hung, L.-W. (2007). Acta Crystallographica Section D 63, 597–610.

Terwilliger, T. C., Ludtke, S. J., Read, R. J., Adams, P. D. & Afonine, P. V. (2020). bioRxiv, 845032.

Terwilliger, T. C., Sobolev, O. V., Afonine, P. V. & Adams, P. D. (2018). Acta Crystallographica Section D 74, 545–559.

Urzhumtsev, A., Afonine, P. V., Lunin, V. Y., Terwilliger, T. C. & Adams, P. D. (2014). Acta Crystallographica Section D 70, 2593–2606.

Wang, B. C. (1985). Methods Enzymol 115, 90–112.

